# A Closed Computational–Experimental Loop Identifies Metabolic Collapse at the Root of Macrophage Dysfunction due to Zinc Dyshomeostasis

**DOI:** 10.64898/2026.01.09.698698

**Authors:** Sunayana Malla, Deandra R. Smith, Ashley M. Peer, Derrick R. Samuelson, Daren L. Knoell, Rajib Saha

**Affiliations:** Department of Chemical and Biomolecular Engineering, University of Nebraska-Lincoln, Lincoln, NE; Department of Pharmacy Practice and Science, College of Pharmacy, University of Nebraska Medical Center, Omaha, NE; Pulmonary Division, Department of Internal Medicine, College of Medicine, University of Nebraska Medical Center, Omaha, NE

## Abstract

Zinc plays a crucial role in immune regulation, the oxidative stress response, and epithelial barrier integrity, yet zinc’s precise role in regulating metabolic and immunological functions in myeloid cells remains poorly understood. Here, we employ a systems biology approach using constraint-based modeling to elucidate the consequences of myeloid-specific loss of ZIP8 on macrophage metabolic function and antibacterial capabilities. We demonstrate that macrophage populations in the lung of ZIP8 knockout (*Zip8*KO) mice exhibit widespread metabolic disruption, spanning glycolysis, butanoate metabolism, amino acid metabolism, and mitochondrial function. Specifically, *Zip8*KO macrophages exhibit impaired nutrient uptake and dysregulated energy metabolism, which is exacerbated following *Streptococcus pneumoniae* infection. Genome-scale metabolic modeling and flux analysis revealed a paradoxical pattern of metabolic suppression prior to infection, followed by overcompensation post-infection, potentially driving immune dysfunction. Consistent with these predictions *Zip8*KO bone marrow-derived macrophages displayed increased ATP demand and disrupted mitochondrial energetics, compromising their ability to control infection. Importantly, we identified succinate, and kynurenic acid as metabolites capable of restoring immune responses and validated their ability to enhance bacterial clearance in *Zip8*KO BMDMs. Together, these findings establish ZIP8 as a central regulator of immune-metabolic homeostasis and suggest potential therapeutic avenues to restore immune function in settings of zinc deficiency.

## INTRODUCTION

Community-acquired pneumonia (CAP) remains a significant global health burden, with *Streptococcus pneumoniae* (pneumococcus) being the primary bacterial cause in the United States^1^. Individuals with compromised immune systems are particularly vulnerable to CAP, with infections often resulting in high rates of hospitalization and mortality^1^. A robust immune defense relies on adequate nutritional support, including essential micronutrients such as zinc (Zn)^2^. Zn is a critical cofactor in numerous biological processes, and its deficiency results in immune dysfunction and increased susceptibility and severity of infections, including pneumonia^3,4^. Globally, an estimated 17% of the population is at risk of insufficient Zn intake, a statistic that underscores the broader implications of Zn imbalance for public health^4^.

Zn is essential for the proper function of myeloid cells, including but not limited to monocytes, macrophages, and dendritic cells. Insufficient intracellular Zn impairs differentiation, phagocytosis, delays the resolution of infection, and hinders wound healing^5^. At the molecular level, Zn interacts with approximately 20% of the human proteome, playing key roles in enzyme activity, signaling, and structural stability^6^. Two families of Zn transporters tightly control intracellular Zn content: ZIPs (Zrt-, Irt-like proteins) and ZnTs (zinc transporters)^5,7^. Among these, ZIP8 has emerged as a key conductor that helps orchestrate and balance the innate immune response against multiple pathogens, including *S. pneumoniae*^7–10^. Given its dual role in Zn transport and immune modulation, our group has previously shown that disruption of ZIP8 expression and function has far-reaching effects on host defense, particularly in the lungs, where Zn biodistribution is essential at the onset of infection^5,7^. We previously reported that in the setting of dietary Zn deficiency and sepsis, mice show increased bacterial burden and mortality despite increased recruitment of myeloid cells to the area of infection^11^. This suggests that the lung macrophage landscape dynamically changes in the absence of ZIP8 and Zn, adversely impacting the ability of these essential phagocytic cells to effectively clear pathogens following lung infection^11^. Given the apparent broad impact that Zn has on shaping the lung myeloid landscape, we sought to understand whether alterations in lung macrophage metabolism may account, in part, for the adverse functional changes observed in response to infection.

To this end, we conducted a comprehensive study to examine potential metabolic shifts as a consequence of ZIP8 loss in macrophages using multiple Genome-Scale Metabolic (GSM) models. A genome-scale metabolic (GSM) model is a comprehensive representation of a cell’s metabolism, encompassing all known metabolic reactions, their associated genes, and metabolites^12^. These models serve as powerful predictive computational tools for studying dynamic biological conditions, including diseases and health-related phenomena, by simulating cellular metabolism under different genetic and environmental contexts^13^. By enabling the simultaneous investigation of multiple metabolic networks in response to perturbations, such as gene deletion or environmental stressors, GSM models facilitate the identification of potential pathogenic mechanisms, biomarkers, and novel therapeutic targets. These insights might not be accessible through traditional experimental or statistical methods alone^12^ without the help of GSM models. For example, gene expression data can reveal which enzymes *Mycobacterium tuberculosis* produces within host macrophages, but GSMs were able to predict that the pathogen adapts by utilizing host-derived cholesterol and fatty acids for survival, a finding later validated experimentally and now explored as a therapeutic target^14–16^. Similar breakthroughs have been achieved in characterizing the metabolic reprogramming of cancer cells, elucidating host-pathogen metabolic interactions in bacterial and viral infections advancing our understanding of immunometabolism^17–19^. These cases highlight the unique ability of GSMs to model and simulate the dynamic, condition-specific behavior of metabolism, offering mechanistic and actionable insights that static molecular data cannot capture^20^.

In this study, we developed macrophage-specific, genome-scale metabolic (GSM) models for both normal wild-type (WT) and ZIP8 knockout (*Zip8*KO) mice before and after *Streptococcus pneumoniae* lung infection, which utilized our single-cell RNA sequencing (scRNA-seq) database to characterize macrophage clusters (**Figure 1**). Myeloid cells, including macrophages, rely on ZIP8 at the onset of infection to generate an effective and balanced innate immune response. This study reveals for the first time that defects in Zn homeostasis disrupt innate immune function in macrophages due, in part, to alterations in several vital metabolic pathways, including central carbon, amino acid, butanoate, fatty acid, lipid, and cholesterol metabolism. Flux analysis attributed all metabolic disruptions to shifts in glycolytic and mitochondrial function, generating a highly unstable environment with flawed amino acid utilization (such as serine, leucine, and alanine) and elevated ROS production. These predictions were validated via Seahorse analysis, which demonstrated a distinct shift in mitochondrial and glycolytic activity in bone marrow-derived macrophages (BMDMs) generated from *Zip8*KO mice before and after infection compared to WT BMDMs. Building on these observations, we applied further optimization techniques, including shadow price analysis and our recently developed MetaShiftOptimizer platform, to identify host-derived metabolic candidate molecules that can restore the ability of ZIP8-defective macrophages to clear engulfed bacteria. *In silico* simulations predicted succinate, N-acetylcysteine (NAC), kynurenine, and 4-aminobutyrate as leading candidates. Strikingly, *in vitro* treatment with succinate and kynurenine before infection led to a significant reduction in bacterial clearance, while NAC, and 4-aminobutyrate had no beneficial effect. This highlights the predictive value of GSMs but also their inherent limitations, as they fail to consider metabolic-signaling crosstalk and immune regulatory feedback. Together, these findings reveal for the first time an essential contribution of metabolic dysfunction in driving immune impairment in ZIP8 deficiency and demonstrate the potential of GSM-based approaches to uncover biologically actionable therapeutic strategies.

**Figure 1.**
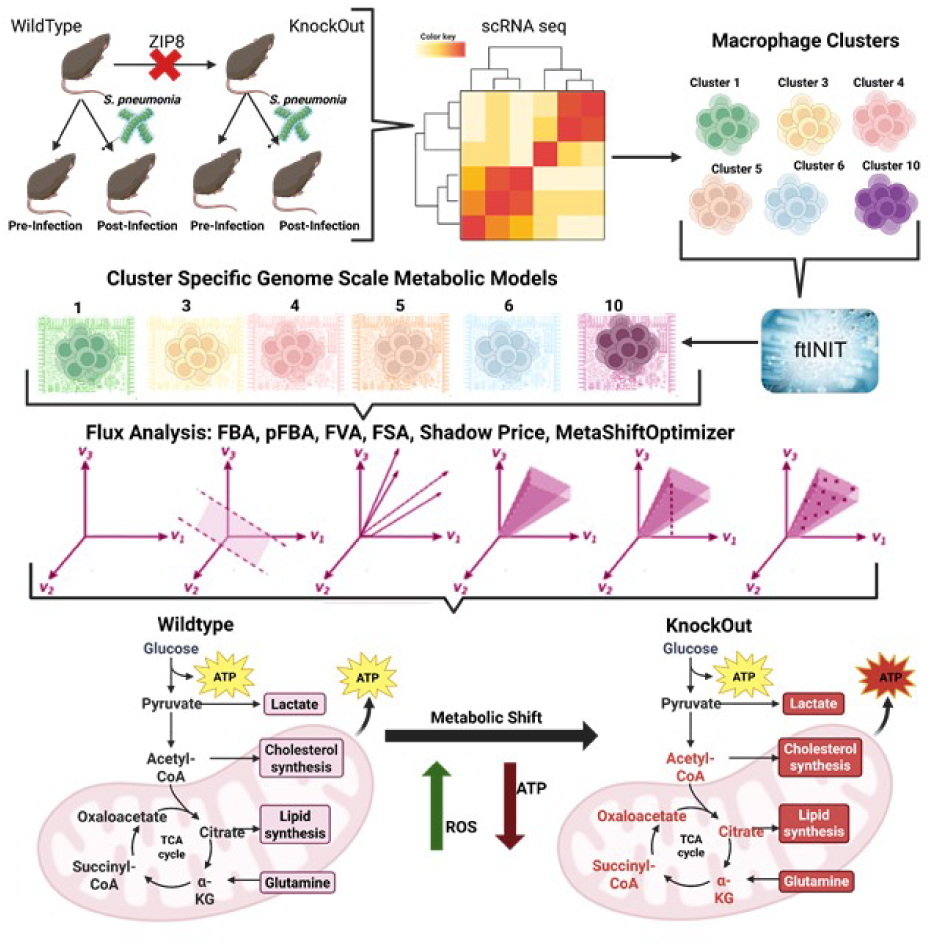
Schematic Work-Flow Model. Starting from the top left and working across and down, scRNA-seq analysis was conducted on lung tissue obtained from WT and *Zip8*KO mouse lung tissue before and after infection with *S.pneumoniae*. Macrophage-specific clusters were identified and then subject to genome-scale metabolic modeling via the ftINIT pipeline. These models were analyzed to identify metabolic capacities and uncover significant metabolic vulnerabilities. This revealed thatZIP8 loss was strongly associated with mitochondrial dysfunction which is associated with disruption of multiple essential metabolic processes, all of which have the potential to impair macrophage-mediated bacterial clearance.

## RESULTS

### Differential Gene Expression and Flux Comparisons Identify Dysregulated Metabolic Pathways as a Consequence of ZIP8 Loss

We previously revealed that ZIP8 loss significantly alters the number and phenotypes of macrophage subpopulations in lungs in response to infection (*manuscript under revision*). Based on this, distinct macrophage clusters were identified by performing graph-based clustering on the scRNA seq datasets ^11^. Next, we comprehensively evaluated the mechanistic consequences of ZIP8 loss in macrophages using exhaustive comparative analysis between the WT and *Zip8*KO conditions with emphasis on macrophage-specific clusters. Prior studies found that *Zip8*KO mice exhibit profound dysregulation in key intracellular signaling cascades that control immune function and bacterial clearance^11,21^. However, a critical question remained as to whether these perturbations are confined to signaling networks or if they extend to metabolic dysregulation at the pathway level. To address this, we performed a differential gene expression (DEG) analysis specifically focusing on metabolism-associated genes^22^. Mapping the differentially expressed genes to metabolic pathways revealed significant perturbations in bile acid synthesis, ascorbate metabolism, and butanoate metabolism, suggesting systemic metabolic remodeling at baseline and post-infection in Z*ip8*KO macrophage clusters compared to noninfected and infected wild-type samples.

Additionally, we leveraged genome-scale metabolic (GSM) models of the identified macrophage clusters (clusters 1, 3, 4, 5, 6, and 10) to compare the metabolic landscapes of WT and *Zip8*KO macrophages pre- and post-infection. Using metabolic models derived from the scRNA seq dataset, we investigated metabolic shifts in WT and Z*ip8*KO macrophage clusters under both pre- and post-infection conditions (**Figure 2a**). To assess the flexibility of metabolic networks in these models, we employed Flux Variability Analysis (FVA), which quantifies the range of allowable flux distributions while maintaining steady state feasibility^23^. FVA thus provides a measure of metabolic plasticity, revealing how individual reaction fluxes may vary within the constraints of the system^13^. We found major disparities in carbohydrate metabolism pathways, including glycolysis, pyruvate, pentose phosphate, galactose, and fructose metabolism. Further, FVA predicted potentially significant adverse changes in oxidative phosphorylation (OxPhos) and reactive oxygen species (ROS) detoxification in *Zip8*KO metabolic models when compared to WT counterparts, in addition to alterations in the majority of amino acid metabolic pathways (**Figure 2b**). These flux patterns were a prominent indication of major metabolic disruption in lung macrophages as a consequence of ZIP8 loss. This raised the question whether these metabolic shifts could result in defects in the ability of macrophages to kill phagocytosed bacteria. Indeed, flux distribution identified a shift in the metabolic landscape at baseline, indicating that the lung macrophages of Z*ip8*KO mice possess significant metabolic derangement prior to infection, and then overcompensate for these changes in response to infection as evidenced by high pathway fluxes in *Zip8*KO post-infection models (**Figure 2b**). We observed this distinct shift in prominent pathways such as glycolysis, OxPhos, valine, leucine, isoleucine, arginine, proline, and lipid metabolism (Fig. 2b).

**Figure 2.**
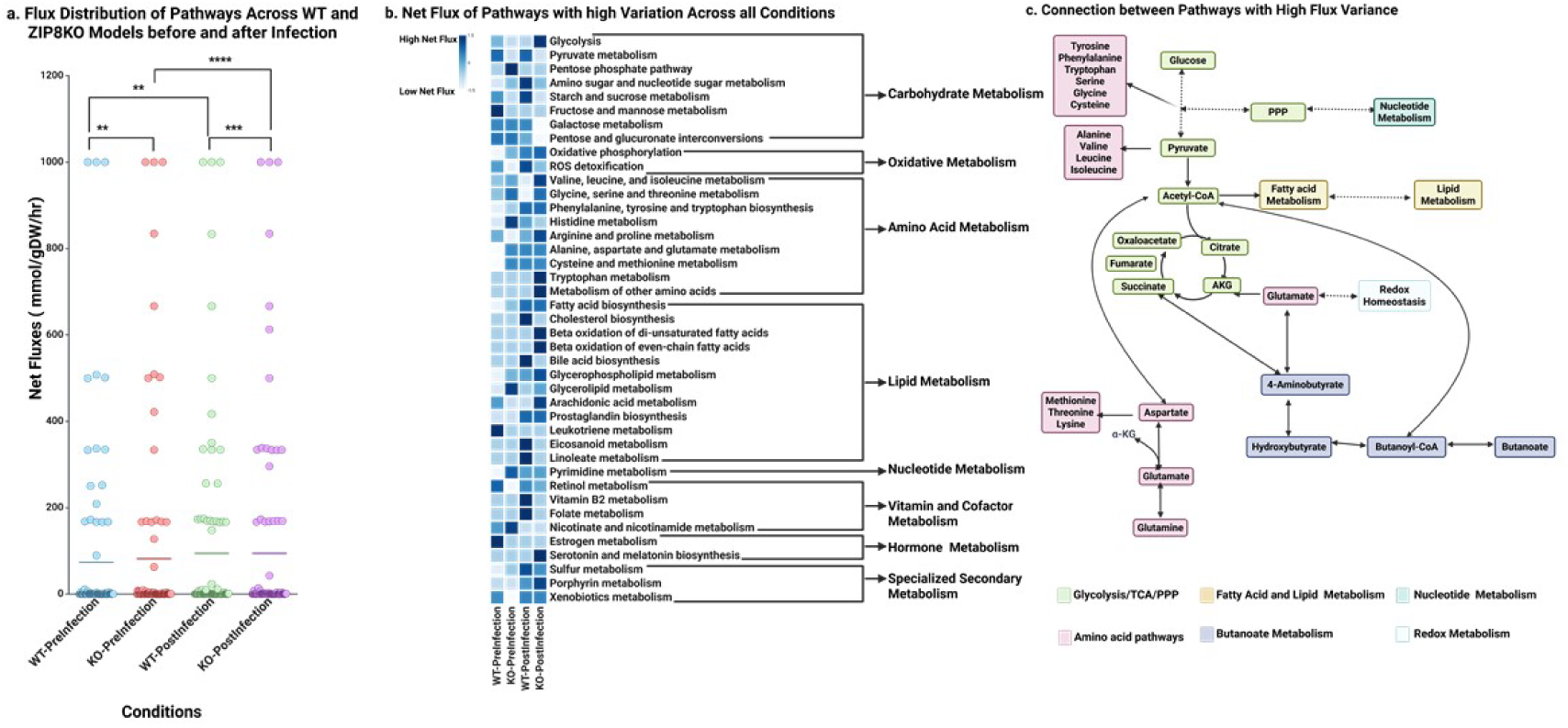
Disrupted Pathways in *Zip8*KO Models Pre-and Post-Infection. (**a**) Scatterplot for pathway net flux distribution in WT and *Zip8*KO macrophage clusters both pre-and post infection show distinct pattern shifts. Variance magnitude was determined by calculating the fold-change of each comparison relative to the smallest observed variance (** FC >= 1, *** FC >= 1.1, and **** FC>= 1.3). (**b & c**) Heatmap and metabolic map of metabolic pathways that vary the most across all conditions-highlighting major shifts originating from all parts of metabolism.

Using the flux data from GSM models combined with DEG analysis, we constructed a metabolic map to explore interconnections between dysregulated metabolic pathways (**Figure 2c**). Given that the pathways identified play key roles in maintaining cellular homeostasis, growth, and immune function, we hypothesized that they may all contribute centrally to the predicted dysfunction. Among these, butanoate metabolism emerged as a key node, not simply due to changes within the pathway itself, but because of its extensive crosstalk with other metabolic systems^24,25^. Specifically, it interfaces with central carbon metabolism (including glycolysis, the TCA cycle, oxidative phosphorylation, and pyruvate metabolism), lipid biosynthesis, and amino acid metabolism (e.g., glycine, serine, threonine, phenylalanine, tyrosine, and tryptophan) (**Figure 2c**). This integrative positioning suggests that disruptions in butanoate metabolism may propagate across multiple networks, amplifying metabolic imbalance and contributing to the overall dysfunction observed in *Zip8*KO mice.

Collectively, our findings reveal for the first time that ZIP8 may have a vast impact on macrophage function through carbohydrate, oxidative, amino acid, lipid, and butanoate metabolism, which are all adversely impacted as a result of ZIP8 loss and subsequent defects intracellular metal transport. The impaired metabolic plasticity revealed by *in silico* analysis suggests that ZIP8 loss disrupts steady-state metabolic flux, as well as innate immune host defense in response to pathogen invasion.

### ZIP8 Loss Impairs Nutrient Utilization and Amino Acid Metabolism

Amino acids not only serve as building blocks for protein synthesis but also function as critical regulators of immune cell signaling, redox balance, and energy metabolism^26^. Given their metabolic interconnectivity and functional significance, we explored how ZIP8 loss influences amino acid utilization and distribution across different cellular compartments. Using a combinatorial approach with Flux Balance Analysis (FBA), parsimonious Flux Balance Analysis (pFBA), and Flux Variability Analysis (FVA), we systematically characterized changes in amino acid transport, synthesis, and degradation. These analyses revealed that organelle (commonly referred as compartment in the modeling context)-specific metabolic alterations may contribute to the reduced metabolic flexibility and impaired immune function observed in *Zip8*KO macrophages.

Flux analysis demonstrated that *Zip8*KO macrophages have reduced amino acid utilization relative to WT counterparts before infection, once again suggesting that metabolic dysregulation precedes pathogen exposure. Following infection, however, *Zip8*KO macrophage clusters overcompensate in amino acid utilization (excluding isoleucine, arginine, and tyrosine), exceeding levels observed in uninfected and infected WT models (**Figure 3a)**. This is remarkable considering that 8–12-week-old adult *Zip8*KO male and female mice appear normal *in vivo*, similar to WT counterparts, despite significant underlying disruption in multiple major metabolic pathways.

**Figure 3.**
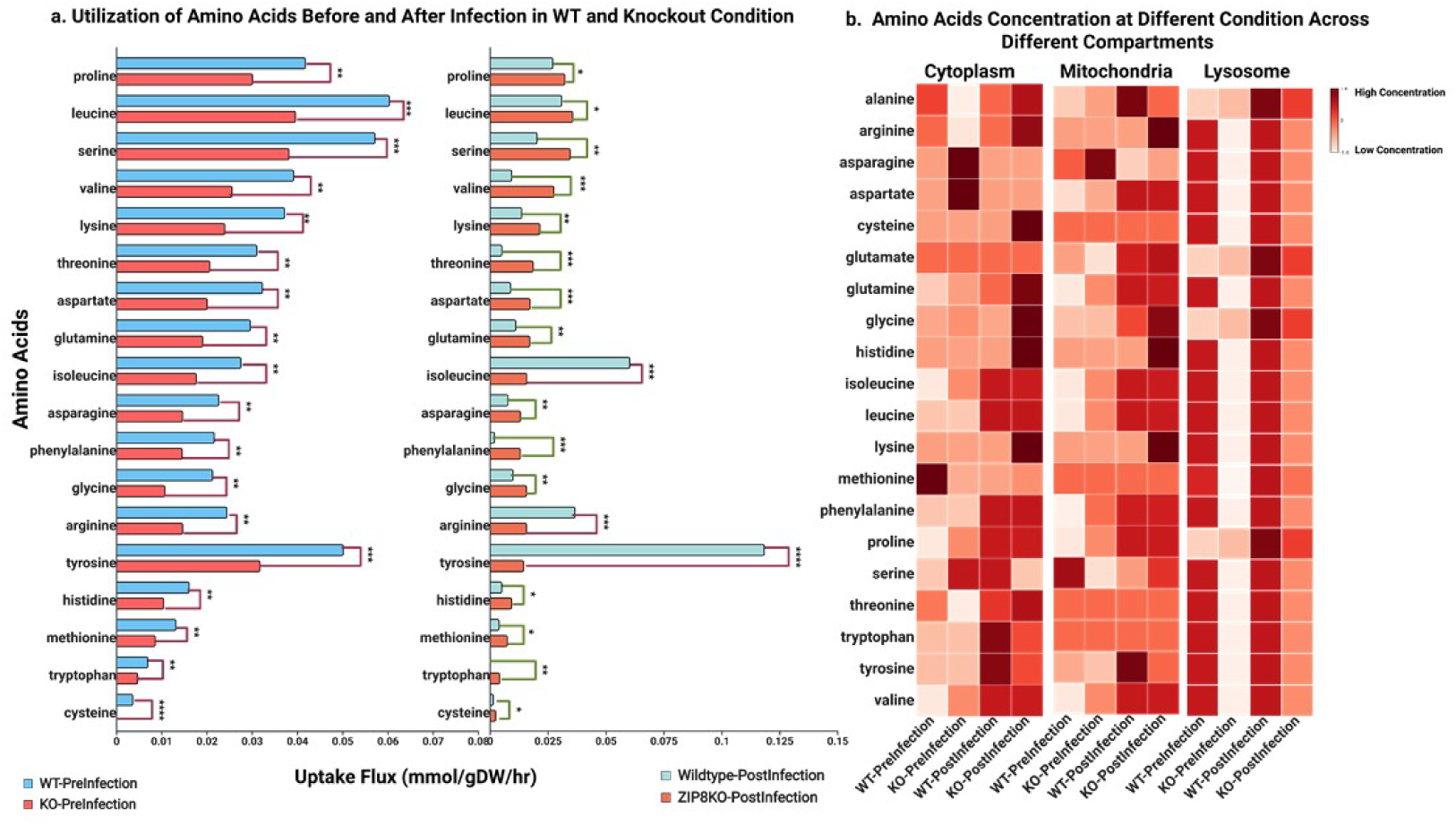
*In silico* Prediction Indicates Major Dysregulation in Amino Acid Metabolism. (**a**) Comparison of amino acid utilization in WT and *Zip8*KO macrophage clusters before and after infection. Significant utilization shift was measured by fold­change of fluxes (unit: mmol/gDW/hr) **FC >= 0.5, *** FC>=l, **** FC>= 5. (**b**) Heatmap showing the Flux Sum of amino acids across three organelle compartments (cytosol, mitochondria, and lysosome). The shift in amino acid utilization and *in silico* concentration indicates major metabolic shifts occurring in *Zip8*KO macrophage clusters.

Previous flux analyses indicated substantial divergence in amino acid-related activity between the WT and *Zip8*KO clusters. To explore this in more detail, we examined *in silico* amino acid concentrations across multiple cellular compartments, including the cytoplasm, mitochondria, and lysosomes. In the noninfected *Zip8*KO model, cytoplasmic amino acid levels exhibited a pronounced imbalance, while glycine and glutamine accumulated to levels significantly higher than those in WT macrophage clusters. In contrast, basic amino acids, including arginine, lysine, and histidine were depleted. This pattern suggests that ZIP8 may play a role in regulating the uptake, synthesis, or intracellular transport of basic amino acids, and that its loss disrupts this balance, potentially leading to cytoplasmic accumulation of other amino acids. A similar trend was observed in the mitochondria, where basic amino acids remained depleted, while glutamine and proline were elevated. These compartment-specific disruptions suggest defects in transport or metabolic regulation, potentially involving mitochondrial and cytoplasmic amino acid handling pathways (**Figure 3b**).

Following infection, divergence between WT and *Zip8*KO models became even more pronounced. While WT macrophages exhibited relatively modest shifts in amino acid levels, representative of a coordinated metabolic and immune response, *Zip8*KO clusters responded with highly dysregulated changes. The amount of cytoplasmic and mitochondrial basic amino acids, glycine, and cysteine significantly increased in *Zip8*KO macrophages, far exceeding what was observed in WT models. The excessive accumulation of glycine and cysteine post-infection is particularly notable, as these amino acids are closely linked to redox homeostasis. This also aligns with flux analysis, which predicted disruption of ROS detoxification pathways in *Zip8*KO models, further suggesting that ZIP8 has a central role in coordinating amino acid metabolism in response to bacterial infection.

In addition to amino acid imbalance, *in silico* modeling predicted a deficiency in butyrate levels in *Zip8*KO macrophage clusters, which coincided with dysregulation of key intermediates in the butanoate metabolic pathway (*manuscript under revision*). Notably, intermediates including 4-aminobutanoate, hydroxybutanoate, and butyryl-CoA, many of which localize in the mitochondria, were significantly altered. These metabolites interface with glutamate, serine, leucine, and fatty acid metabolism, indicating a broader disruption of interconnected metabolic networks. Together, these observations suggest a biphasic metabolic trajectory in *Zip8*KO macrophage clusters, involving an initial deficiency in key metabolites and metabolic activity, likely due to disrupted Zn homeostasis via impaired ZIP8 transport, which then manifests as an acute hyperactive response to infection characterized by metabolite accumulation. We contend that this dysfunctional metabolic response likely contributes to the exaggerated proinflammatory, and destructive phenotype observed in the lungs of *Zip8*KO mice in response to *S. pneumoniae*^11^.

The widespread alterations across multiple amino acid pathways, alongside disruptions in butanoate, fatty acid, bile acid, and cholesterol metabolism, led us to question whether a central master regulatory hub is responsible for many of these diverse imbalances. Accordingly, we next evaluated whether there exist major shifts in carbohydrate and oxidative phosphorylation metabolic pathways with emphasis on glycolysis, the pentose phosphate pathway (PPP), oxidative phosphorylation (OxPhos), and ROS detoxification.

### ZIP8 Loss Disrupts Glycolytic and Mitochondrial Function in Macrophages and Inhibits Bacterial Clearance

Computational analysis revealed substantial alterations in amino acid metabolism, impaired nutrient uptake, and disrupted intermediates in butanoate metabolism. These findings led us to determine what factors drive the underlying metabolic reprogramming in *Zip8*KO macrophage clusters. Notably, flux analysis revealed a marked increase in ATP utilization in *Zip8*KO macrophage clusters in response to infection, with ATP demand and consumption the highest in *Zip8*KO clusters post-infection compared to WT pre-infection, *Zip8*KO pre-infection, and WT post-infection (**Figure 4a**). ATP availability is critical for nutrient uptake and transport, including amino acids. The shift observed in ATP demand in the *Zip8*KO macrophages in response to infection indicates that a major bottleneck in these processes is likely to significantly contribute to changes in glycolytic and fatty acid metabolic activity, as well as key amino acid pathways. Collectively, these observations also strongly imply that mitochondrial function is significantly dysregulated as a result of ZIP8 loss, which may also contribute to defective macrophage-mediated clearance of bacteria. Based on this, we next determined whether any host-derived molecules, or lack thereof, may be driving changes in metabolism.

**Figure 4.**
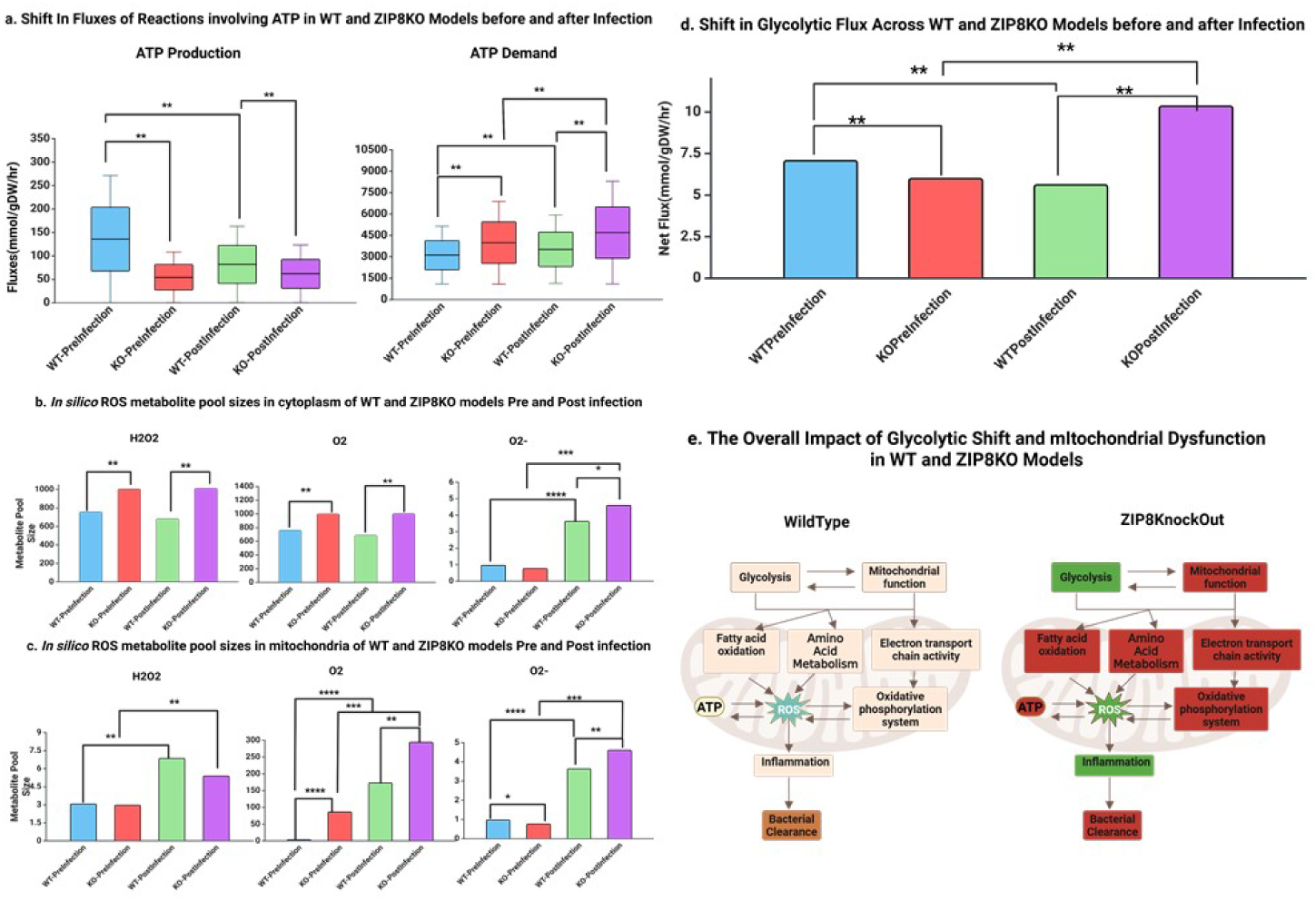
Glycolytic and Mitochondrial Dysfunction as Drivers of *Zip8*KO Dysregulation. (**a**) Box plot comparison of ATP production and demand in WT and *Zip8*KO macrophage clusters before and after infection. (**b, c**) Comparison of differences in the predicted pool size of reactive species and oxygen in the cytoplasm and mitochondria of WT and *Zip8*KO macrophage clusters. (**d**) Changes in net glycolytic flux (mmol/gDW/hr) in WT versus *Zip8*KO macrophage clusters. (**e**) Schematic diagram connecting the potential for metabolic changes to influence immune function. Significant change influxes is calculated by Fold Change (FC) where ** FC >= 0.5, *** FC >= 3, **** FC >= 5.

We found that increased ATP demand in *Zip8*KO macrophage clusters after infection was accompanied by significant reductions in succinate and ubiquinone levels, which were also reduced in *Zip8*KO clusters before infection when compared to WT models. Amino acid and butyrate fluxes were also upregulated post-infection, whereas most *Zip8*KO clusters exhibited a decrease prior to infection. These findings strongly indicate metabolic imbalance (**Figure 4b and 4c**), where *Zip8*KO macrophage clusters fail to efficiently meet ATP demands, leading to disrupted energy metabolism that impairs multiple interconnected pathways. The drastic increase in ATP consumption in response to infection also suggests that the system is attempting to compensate for pre-existing deficiencies but may ultimately fail to sustain metabolic homeostasis, potentially contributing to an ineffective immune response. ATP-dependent transport processes are particularly crucial for amino acid metabolism, fatty acid oxidation, and butanoate metabolism, all of which require high energy input for processing and assimilation^27,28^. A shift in energy metabolism would therefore likely disrupt central metabolic pathways, causing macrophage dysfunction in response to pathogens.

To further explore the role of oxidant imbalance, we used *in silico* metabolite pool sizes of reactive oxygen species (ROS) to define the extent of mitochondrial dysfunction. Consistent with oxidant imbalance, the relative amount of cytosolic ROS species, including H_2_O_2_, and O_2_-increased in *Zip8*KO macrophages (**Figure 4b & 4c**). The increase in cytosolic ROS indicates that macrophages encounter oxidative stress, which is further amplified due to concomitant defects in the ROS detoxification pathway. In comparison, we observed a significant increase in O_2_ and O_2_-in the mitochondrial compartment, especially following infection, when compared to WT macrophage clusters. The change in ROS metabolite pool sizes in tandem with the changes in ATP demand and amino acid metabolism strongly suggests that mitochondrial dysfunction is a root cause of defective immune function and tissue disrepair because of ZIP8 loss in macrophages. Notably, we also observed a significant shift in glycolysis, characterized by increased glycolytic flux in *Zip8*KO models compared to WT controls **(Figure 4d**). Enhanced glycolytic activity is consistent with the metabolic profile of proinflammatory macrophages and aligns with the proinflammatory gene expression profiles in macrophage clusters showing that most macrophages in the lung of infected *Zip8*KO mice are of a pro-inflammatory phenotype. We also observed an increase in PPP activity in *Zip8*KO clusters before infection, which declined precipitously following infection. One potential interpretation of this finding is that elevated glycolysis and PPP activity in the *Zip8*KO mouse lung before infection indicates once again that metabolic derangement exists at baseline and contributes to defective innate immune function and the ability to eradicate bacteria. The decrease in PPP activity post-infection also suggests that the *Zip8*KO models are not able to properly fight infection^11^. These findings further underscore the critical role of ZIP8 in central carbon metabolic function, including the glycolytic, pentose phosphate pathways, and energy metabolism (**Figure 4e**).

To validate *in silico* metabolic predictions, we assessed mitochondrial respiration of both *Zip8*KO and WT BMDMs before and following infection with *S. pneumoniae* using the Seahorse XFp Mito Stress Test (**Figure 5a**). *Zip8*KO macrophages displayed a higher oxygen consumption rate (OCR) compared to WT BMDMs at baseline (**Figure 5b**), which was further exacerbated in response to infection (**Figure 5c**). Following oligomycin injection, both uninfected and infected *Zip8*KO macrophages displayed increased ATP-dependent cellular respiration compared to WT, as well as significantly higher maximal respiration following administration of the mitochondrial uncoupler FCCP (**Figure 5d**). Consistent with this, we observed a similar and significant increase in Mitosox Red staining of *Zip8*KO BMDMs, indicative of increased ROS generation, at baseline and following infection when compared to WT BMDMs (**Figure 5e, 5f, & 5g**). We also assessed the extracellular acidification rate (ECAR) using the Seahorse Glycolysis stress test. Consistent with computational metabolic predictions, we observed that *Zip8*KO BMDMs rely more heavily on glycolysis as a source for ATP synthesis compared to WT BMDMs (**Figure 5h & 5i**). Furthermore, the glycolytic reserve of both *Zip8*KO and WT BMDMs, independent of infection, is nonexistent, suggesting that these cells have already reached their glycolytic limit (**Figure 5g & 5h**). Together, these data confirm that *Zip8*KO BMDMs are metabolically compromised prior to infection through upregulation of oxidative and glycolytic pathways to accommodate increased energy demand. Collectively, these findings validate computational predictions and demonstrate that ZIP8 loss alone increases cellular ATP demand, which then compromises energy utilization in response to pathogen engulfment, contributing to an inability to efficiently clear phagocytosed bacteria.

**Figure 5.**
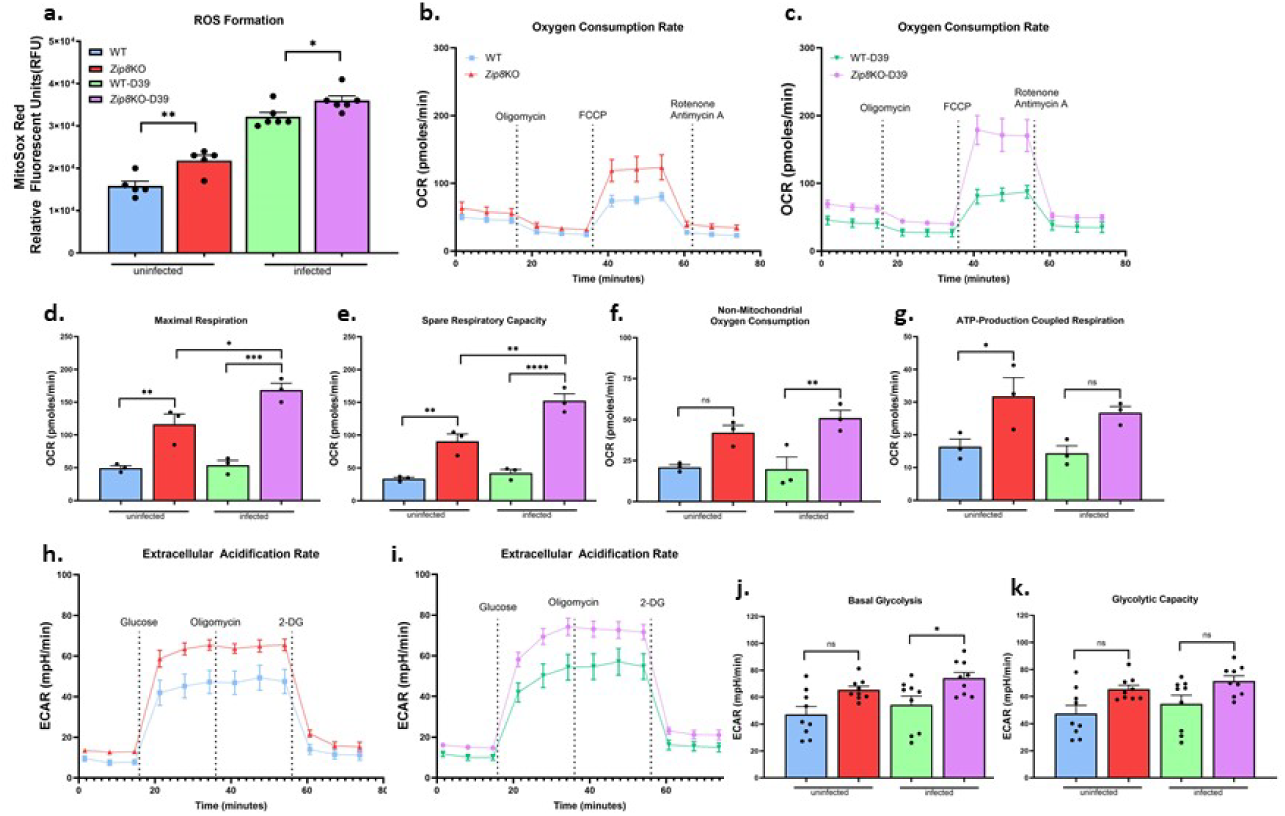
Validation of Predicted Metabolic Dysfunction in *Zip8*KO BMDMs Before and After Infection. WT and *Zip8*KO BMDMs were analyzed before and after *S.pneumoniae* infection for: **(a)** Intracellular ROS as determined by MitoSox staining; **(b-g)** the SeaHorse XF Mito Stress test; and **(h-k)** the XF Glycolysis Stress Test. As predicted by *in silico* analysis, *Zip8*KO BDMSs have significantly increased ROS production, oxygen demand, extracellular acidification, and glycolytic activity compared to WT BMDMs that is further exacerbated in response to infection. (*p < 0.05; **p < 0.01; ***p <0.001. Data representative of a minimum of 3 independent experiments; Data were analyzed using GraphPad Prism version 10.5.0. Statistical significance was determined using one-way ANOVA followed by Sidak’s corrections for multiple comparisons between groups. Values are expressed as means + SEM).

### *In silico* Identification of Metabolic Intermediates that Restore Bacterial Clearance in the Setting of Zinc Dyshomeostasis

Next, we conducted an integrative analysis of macrophage clusters in all models to identify deficits in host-derived metabolic byproducts that would also be predicted to have therapeutic potential to restore bacterial clearance. Knowing that systems-level assessment revealed profound disruption of energy metabolism in *Zip8*KO BMDMs, we interrogated metabolic pathways involved in glycolysis, mitochondrial function, and amino acid, fatty acid, and bile acid metabolism to identify leading molecules. To systematically evaluate these perturbations, we employed a suite of computational approaches that included shadow price analysis, which identifies metabolites that exert the highest control over metabolic flux distributions^29^. This method quantifies the sensitivity of metabolic networks to changes in metabolite availability and thereby reveals candidate molecules whose reincorporation into the model could reestablish coherence between predicted fluxes and transcriptomic profiles. Comparative shadow price analysis revealed aberrant regulation of key mitochondrial metabolites, including succinate, fumarate, and glutamate. These findings informed a refined list of candidate metabolites with the potential to rescue defective immune responses by targeting mitochondrial and auxiliary metabolic pathways. Additionally, we utilized MetaShiftOptimizer^30^, a bilevel optimization framework that we recently developed, to pinpoint reactions that could phenotypically shift *Zip8*KO BMDMs back toward WT (normal) behavior^30^. This identified specific reactions whose upregulation, downregulation, or deletion could align knockout metabolic fluxes with those of WT macrophages. Using combined computational approaches, we identified 4-aminobutanoate, succinate, N-acetylcysteine (NAC), and kynurenine as leading compounds with potential to restore bacterial clearance in *Zip8*KO BMDMs. To assess their impact, we also compared the metabolic flux profiles to the *Zip8*KO pre-infection model before and after the addition of each compound further substantiating that 4-aminobutanoate, succinate, N-acetylcysteine (NAC), and kynurenine induce significant flux shifts *in silico*, suggesting their potential to restore immune function (**Figure 6a**). Next, we determined whether *in silico* intermediates can restore bacterial clearance following infection of *Zip8*KO BMDMs. From a translational perspective, knowing that ZIP8 variant alleles can prospectively be identified across populations, we utilized a preventive treatment strategy. Briefly, we first conducted dose escalation studies to determine an optimal dose range without toxicity (data not shown). *Zip8*KO and WT BMDMs were then incubated with each compound individually before infection with *S. pneumoniae*. Following overnight incubation, cells were lysed, plated, and bacterial CFUs were enumerated. Strikingly, kynurenic acid (**Figure 6b**) and succinate (**Fig. 6c**) restored the ability of *Zip8*KO BMDMs to eradicate internalized bacteria, whereas neither compound had an impact on infected WT cultures. In contrast, pretreatment with N-acetyl-cysteine (**Figure 6d)** and 4-aminobutyrate (**Figure 6e**) did not improve bacterial clearance. Importantly, all the compounds tested did not affect bacterial viability, as bacteria incubated with each compound at the highest concentration exhibited no differences in *S. pneumoniae* viability following incubation (data not shown).

**Figure 6.**
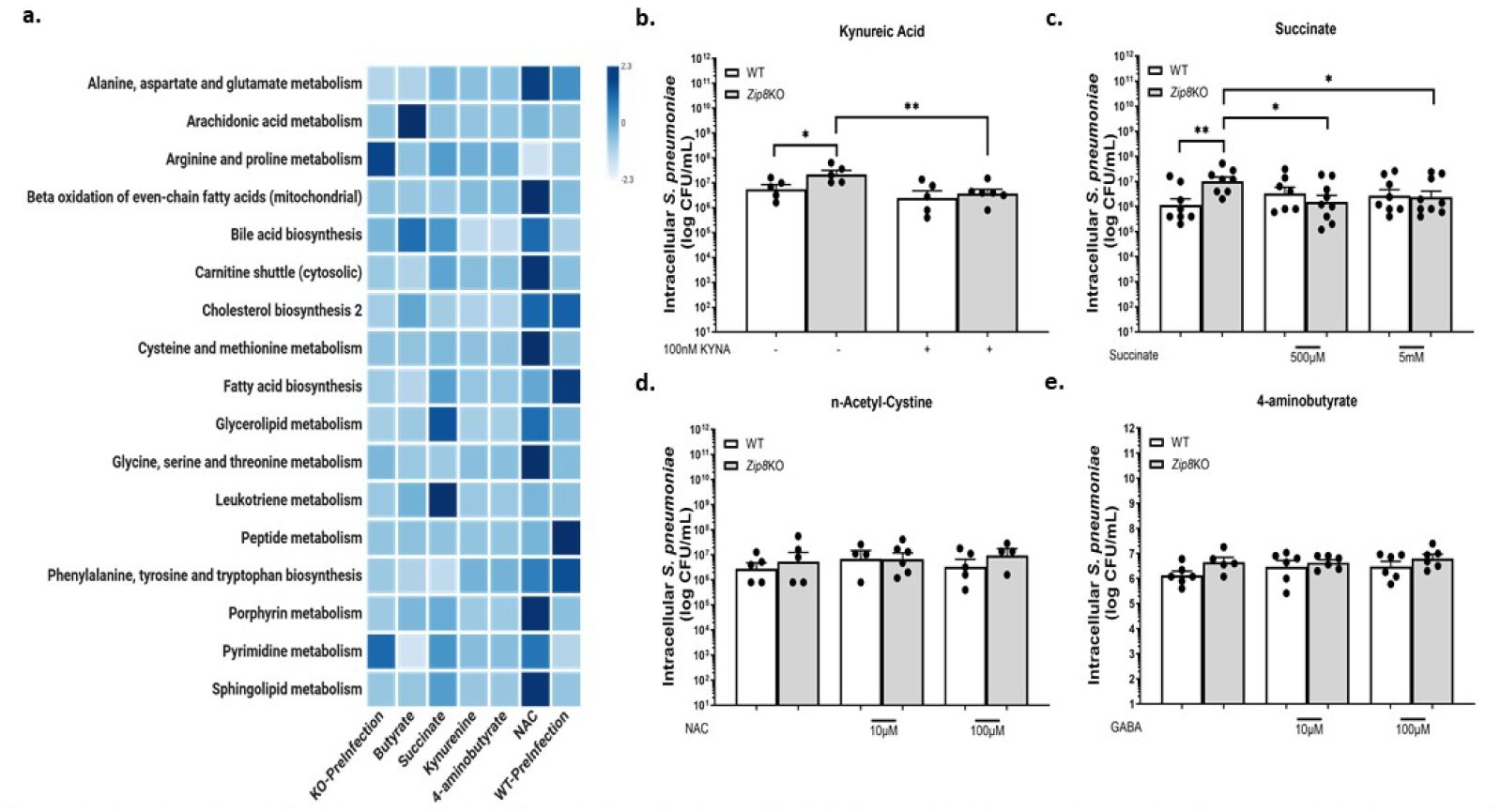
*In silico* Identification and Validation of Host-Derived Metabolites that Restore Bacterial Clearance. (**a**) Heatmap of leading *in silico*-host-derived metabolites via Flux shift analysis reveals kynureic acid, succinate, n-Acetyl-Cysteine, and 4-aminobutyrate as lead candidates. Pretreatment of WT and *Zip8*KO BMDMs with **(b)** kynureic acid and **(c)** succinate restored bacterial clearance in *Zip8*KO BMDMs similar to WT controls. Pretreatment with **(d)** N-acetyl-cysteine and **(e)** 4-aminobutγrate had no beneficial effect. (* p < 0.05; ** p <0.01; Data representative of a minimum of 3 independent experiments; Data were analyzed using GraphPad Prism version 10.5.0. Statistical significance was determined using an unpaired t-test. Values are expressed as means ± SEM).

### Comparison of Metabolic and Signal Transduction Pathways in Wildtype vs Knockout Conditions

Intracellular signaling pathways are often regarded as the primary drivers of immune regulation. Our findings indicate that metabolic pathways also play an important and front-line role in coordination with signal transduction pathways, as a root cause of immune dysfunction. In fact, we contend it is biologically unlikely that only one system can be compromised without impacting the other, whether it be before or following insult. *In silico* analysis revealed profound disruptions in nutrient uptake and energy metabolism in the setting of ZIP8 loss in lung macrophage populations. Given the vital role of mTORC1 in nutrient sensing, ATP regulation, lysosomal biogenesis, and bacterial clearance, we examined whether this signaling pathway could be involved in immune dysfunction in the setting of Zn dyshomeostasis. To this end, we constructed a mechanistic model positioning metabolic shifts as the upstream drivers of immune dysregulation in *Zip8*KO macrophages (**Figure 7**). As anticipated, at baseline, WT clusters exhibited optimal concentrations of key amino acids across intracellular compartments essential for maintaining proper signaling activity. Specifically, mitochondrial integrity ensures appropriate utilization and trafficking of amino acids, including serine, alanine, glutamate, and leucine^31–33^. In particular, amino acids participate in activating and modulating signaling pathways, including mTORC1, in a compartmentalized, organelle-specific manner. Coupled to this, ZIP8 loss in macrophages causes mitochondrial dysfunction in part through altered ATP supply and demand, creating an imbalance that results in increased ROS formation accompanied by widespread disruption in amino acid metabolism. Furthermore, in support of this, we observed a organelle-specific reduction of leucine and arginine, both of which are known to directly activate mTORC^32,33^ In summary, our analysis of the sc-RNAseq lung database revealed that mitochondrial perturbations propagate through metabolic networks, altering the biosynthesis and distribution of multiple amino acids that are also vital for proper cell signaling cascades. This cascade of metabolic dysregulation compromises signaling fidelity that impairs macrophage-mediated innate immune function, resulting in defective bacterial clearance, tissue dysfunction, and likely a deficit in the host’s ability to effectively resolve infection and injury.

**Figure 7.**
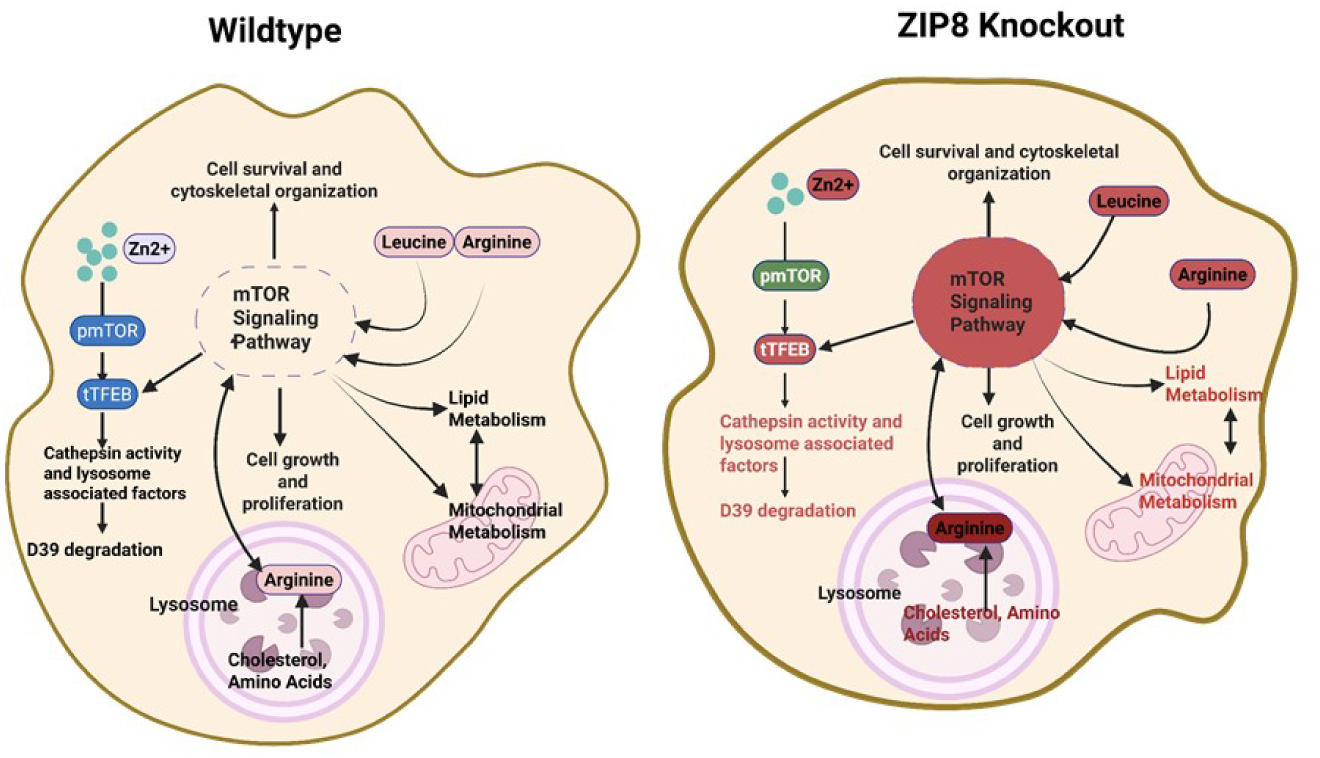
Disruption in Mitochondrial Function Impacts the Overall Physiology, including Signaling Cascades. Based on *in silico* analysis, derangement of multiple metabolomic pathways appeared in *Zip8*KO macrophage clusters compared to WT clusters before and after infection. Many pathways converged directly or indirectly upon the mTOR signaling pathway, a pathway that is vital for proper phagolysomal-mediated removal of bacteria. Red denotes molecules within the *Zip8*KO model whose activity is predicted to be significantly decreased compared to WT models. Green denotes molecules with increased activity.

## DISCUSSION

The integration of diverse flux analysis techniques facilitates a comprehensive understanding of the underlying mechanisms of diseases, thereby enabling the identification of effective therapeutic targets that advance precision medicine and healthcare^34^. In this study, we systematically employed multiple flux analysis techniques, including FBA^35^, FVA^23^, pFBA^36^, FSA^37^, shadow price analysis and MetaShiftOptimizer, to investigate the pathways, reactions, and metabolite flux distributions in lung macrophage models that were derived from our scRNA seq datasets obtained from WT and *Zip8*KO mouse lungs before and following *S. pneumoniae* infection. The results of our study reveal underlying defects in mitochondrial and glycolytic function subsequent to ZIP8 loss and altered intracellular Zn homeostasis. Despite no differences in the overall health and appearance between adult WT and *Zip8*KO mice, there existed underlying major defects in cellular metabolism that we contend created a *“calm before the storm”* situation, resulting in a highly dysfunctional, imbalanced, inflammatory response, which drives excessive collateral tissue damage, decreased bacterial clearance, and decreased survival.

Zn homeostasis is tightly regulated and controlled by ZIPs and ZNTs, where ZIPs are responsible for increasing the cytosolic zinc, whereas ZNTs do the opposite ^38^. ZIP8 is the only ZIP out of 14 whose expression is rapidly induced upon bacterial recognition and required for a balanced and effective macrophage-mediated immune response^11^. It has already been established that ZIP8-mediated Zn mobilization into macrophages and dendritic cells following infection is necessary for dampening the canonical NF-kB signaling pathway and improving recovery from pneumococcal pneumonia ^8^. In related work, we have revealed a similar impact on the mTORC1/TFEB signaling pathway and lysosomal-mediated bacterial clearance of macrophages, whereby ZIP8 loss creates an intracellular Zn deficit that impairs lysosomal biogenesis and function decreasing clearance of phagocytosed bacteria (*manuscript under revision*). The primary focus of this investigation was to determine the extent to which disruption of metabolic pathways determines cell fate in the setting of Zn dyshomeostasis^39^. To achieve this, GSM models were employed to thoroughly explore the metabolic landscape of both WT and *Zip8*KO lung-derived macrophage clusters. GSMs integrate multi-omics data, which include genomics, transcriptomics, and metabolomics, to construct personalized metabolic models that help identify disease-associated metabolic alterations^34^. These models facilitate biomarker discovery by predicting flux differences between healthy and diseased states, aiding in the early detection and prognosis of conditions such as cancer and metabolic disorders^12^. Among several methods available to reconstruct GSM models, ftINIT is particularly beneficial in biomedical research as it enables the development of tissue, cell-type, and condition-specific metabolic models, which are critical for understanding disease mechanisms and drug responses that foster precision medicine^34^. This efficiency is crucial when working with large-scale datasets, including single-cell RNA sequencing and multi-omic studies. Hence, we used ftINIT to reconstruct 24 total GSM models of macrophage clusters (1, 3, 4, 5, 6, and 10) across all *in vivo* conditions (wildtype and *Zip8*KO pre- and post-infection).

Our findings reveal, for the first time, a complex landscape of metabolic and transcriptional disruptions as a result of ZIP8 loss in macrophages that contribute to their impaired immune response to bacteria. The identification of metabolic bottlenecks, particularly in energy supply and demand (glycolysis, PPP, and OxPhos), amino acid metabolism, and butanoate metabolism, highlights a profound reprogramming of cellular metabolism in the absence of ZIP8. The observed shift resulting in reduced oxidative phosphorylation and increased flux towards alternative energy-generating pathways such as glycolysis and PPP, suggests a compensatory adaptation, a phenomenon commonly seen in models of mitochondrial stress and metabolic inflexibility^40,41^. This altered energy state may compromise the cells’ ability to meet the demands of a robust immune response. Importantly, the disruptions extended beyond core metabolism. The impaired nutrient uptake we observed suggests altered cellular homeostasis as a result of Zn dyshomeostasis that adversely impacts production and/or biodistribution of vital metabolic factors, resulting in metabolic stress. We predict that the underlying stress could also restrict the availability of key substrates, amplifying the dysfunction across other biosynthetic pathways that impact immune function^42^. What sets this investigation apart is the emerging evidence that these metabolic changes are not isolated events but are directly coupled to signaling cascades. We identify two interconnected mechanisms of immune impairment in *Zip8*KO macrophages that include a fundamental metabolic deficiency predisposing cells toward a dysfunctional state even before infection and subsequent activation of key signal transduction pathways. This is in part due to a compartmentalized breakdown in amino acid homeostasis in key organelles that is also coupled to the regulation of key signaling pathways. These findings challenge the conventional paradigm that centers signaling cascades as the sole regulators of immune function, instead highlighting metabolism, particularly mitochondrial and amino acid pathways, as a primary determinant of immune competence. This supports growing recognition that metabolism and signaling are deeply interconnected, where a change in one reciprocates a change in the other^43^. In the context of ZIP8, Zn, or lack thereof, we contend that metabolic constraints are also a direct cause for disruptions in immune signaling, oxidative stress responses, and cellular repair mechanisms. If correct, this further challenges the traditional view that signaling alone dictates immune behavior^44^, which also aligns with findings in other disease states, including cancer and chronic inflammation, where metabolic rewiring plays a pivotal role in shaping cell fate^17,45,46^.

To explore the translational relevance of our computational predictions, we tested four metabolomic-based interventions aimed at restoring bacterial clearance in ZIP8-deficient macrophages. Succinate and kynurenic acid, when administered before infection, restored the ability of macrophages to clear internalized bacteria, which is consistent with flux-correcting *in silico* predictions. While these findings are encouraging, additional studies are warranted that dissect the biological impact on associated metabolic and signaling pathways, in addition to preclinical analysis to evaluate efficacy and toxicity. For example, did pretreatment also correct defects in oxidative phosphorylation and glycolysis, and was this required for restoration of bacterial clearance? Whether intracellular deficits in Zn were corrected remains to be determined and whether manganese (Mn++) and iron (Fe++), also ZIP8 substrates, may also influence metabolic behavior^47^. Likewise, NAC and 4-amino-butyrate failed to improve bacterial clearance despite modeling predictions underscoring the complexity of metabolic regulation that constraint-based models may not fully capture^48^. Several factors may underlie this mismatch^49^. First, constraint-based models assume steady-state conditions and optimality principles, which do not always reflect the dynamic, non-equilibrium behavior of real cellular systems^12,50^. Second, enzyme kinetics and substrate affinities are not explicitly modeled, meaning that predicted flux distributions may not translate into actual enzymatic activity under physiological conditions^49^. Third, post-translational modifications, allosteric regulation, and signaling cascades can significantly alter metabolic enzyme activity and immune signaling independent of transcript levels or flux capacity^48^. Fourth, the timing of treatment may require refinement. For example, studies have shown that macrophages exposed to butyrate after differentiation do not have improved immune function; however, butyrate exposure during differentiation enhances macrophage immune function^25^. This indicates that metabolomic-derived preventive treatment strategies may require additional consideration that includes exposure during cellular programming as progenitor cells mature into fully differentiated cells. Fifth, metabolic compartmentalization, particularly between cytosolic and mitochondrial pools, can affect the accessibility and utilization of intermediates like 4-aminobutyrate. The *in vivo* bioavailability of 4-aminobutanoate, succinate, N-acetylcysteine (NAC), and kynurenine differ.

Moving forward, incorporating a layered, multi-omics approach that includes proteomics, metabolomics, and phosphoproteomics data will be crucial for refining model predictions and capturing the full scope of cellular responses to metabolic interventions^12^. Integration with dynamic modeling approaches and machine learning frameworks may also help overcome the limitations of current static models^12,51^. Together, our findings emphasize the importance of viewing immune dysfunction through a metabolic lens. Our findings demonstrate the potential of metabolic modeling to identify pathogenic mechanisms at the cellular and molecular levels to guide therapeutic discovery in a targeted and mechanistically informed manner. While this study provides valuable insights into the metabolic consequences of *S. pneumoniae* infection in the setting of Zn dyshomeostasis, it is important to acknowledge certain limitations. The reconstruction of condition-specific models relies on gene expression data, which may not always directly translate to enzymatic activity^12,50^. Furthermore, metabolic flux predictions assume steady-state conditions, which may not fully capture transient metabolic adaptations occurring in dynamic cellular environments^34^. In the future, with further advancements, GSM models can accommodate additional factors that include thermodynamics and temporally spatial data to further gain deeper insights. Overall, our findings provide a deeper understanding of the metabolic reprogramming associated with ZIP8 loss and Zn dyshomeostasis. When considering that Zn interacts with approximately 20% of the human proteome, our multidynamic computational approach was able to identify metabolic vulnerabilities and therapeutic targets that otherwise may not have been possible. By bridging genome-scale modeling with experimental validation, this study underscores the importance of a systems-level perspective in unraveling disease mechanisms and designing targeted interventions. Further exploration of metabolic plasticity and signaling interactions in similar models may accelerate the discovery of novel therapeutic strategies for metabolic and immune-related disorders.

## STAR METHODS

### Animal Use and Care

At the University of Nebraska Medical Center’s (UNMC) Animal Resource Facility, every animal was kept in pathogen-free environments. Water and food were available at all times. The Institutional Animal Care and Use Committee approved the study protocol employed in these studies. The Office of Laboratory Animal Welfare and the National Institutes of Health provided recommendations for all procedures and methods involving animal care used in this study protocol.

Conditional *Zip8* knockout mice, or *Zip8*KO for short, were produced using the methods previously mentioned^52^. In short, the Neo cassette next to the upstream loxP site was deleted by breeding heterozygous *Zip8^flox-neo/+^* mice to ROSA26:FLPe knock-in mice (The Jackson Laboratory) that expressed FLP1 recombinase ubiquitously. *Zip8^flox/flox^* mice were created by mating the resulting *Zip8^flox/+^* mice. PCR and DNA sequencing validated the loxP sites flanking exon 3 and confirmed the removal of the FRT-flanked region. To create the conditional *Zip8*KO, *Zip8^flox/flox^* mice were bred to LysMcre (The Jackson Laboratory), which is specific to myeloid cells. Jackson Labs provided C57BL/6J wild-type equivalents, which were produced specifically for research purposes. All appropriate measures were taken to ensure proper animal ethics and care using the methods previously described^11,39,53^.

### Single Cell RNA Sequencing, and Macrophage Cluster Identification

Lungs from 1 mouse per treatment group were processed as previously described^54^. Briefly, the lungs were perfused with 10 mL of heparin-PBS. The GentleMACS dissociator (Miltenyi Biotech) was used to separate the harvested lungs from the digesting solution (collagenase I, 0.2 μg/μL + DNase I, 75 U/mL + heparin, 1.5 U/mL) in Dulbecco’s Modified Eagle’s Media (DMEM). Samples were shaken for 30 minutes at 37°C. PBS with 4 mM EDTA was used to counteract the activity of the digestion solution. Red blood cells were lysed for 1 minute in 1 mL of lysis solution (BioLegend), and then neutralized with ice-cold Gibco DMEM. Dead cells were removed using the Dead Cell Removal Kit (Miltenyi Biotec). FACS buffer (2% fetal bovine serum (FBS) + 0.1% NaN_3_ in PBS) was used to prepare cells for RNAseq. All reagents, unless specified, were purchased from Sigma. The viability and quantification of single-cell suspensions obtained from the whole lung were determined using a LUNA-FLTM Dual Fluorescence Cell Counter (Logos Biosystems). Following manufacturer recommendations, single cells were extracted from cell suspensions (100–2,000 cells/μL) using a 10x Chromium controller (10x Genomics). The gel beads emulsion (GEM)/sample solution was collected and placed into strip tubes. Following cDNA amplification, reverse transcription was carried out using a thermocycler (C1000 TouchTM Thermal Cycler, Bio-Rad). Amplified products were solid phase reversible immobilization (SPRI) bead-purified and evaluated by a Fragment Analyzer (Agilent, Santa Clara, CA). Twenty-five percent of the cDNA volume was fragmented, and PCR purification and clean-up were carried out using double-sided SPRIselect (Beckman Coulter). Following adaptor ligation, each sample underwent SPRI clean-up and PCR amplification utilizing sample-specific indexes. The Fragment Analyzer was used to quantify, purify, and determine the library size distribution of PCR products. PCR products were purified, quantified and sequenced on an Illumina (San Diego, CA, USA) Novaseq6000. Libraries were sequenced to an average depth of 50,000 reads per cell following manufacturer-recommended conditions.

### Bioinformatic Analysis

The 10xGenomics cellranger (version 7.1) analysis pipeline was used for demultiplexing and generating a gene-barcode matrix. Sub-function mkfastq from cellranger was used to generate raw read files in FASTQ format; sub-function count was used to map the raw reads to the mouse reference genome (version 10), then performed filtering, cell barcode counting and UMI counting. As a result, a gene expression matrix was generated containing the raw UMI count for each gene for each cell per sample. After generating matrices for all samples, sub-function aggregation was called to aggregate all four samples, all matrices were normalized by subsample reads from higher-depth GEM wells to make them comparable. The aggregated matrix then feed into 10X Loupe Browser for downstream analysis. Cells with more than 5% mitochondrial reads are filtered out. Then applied principal component analysis (PCA) to reduce the dimensions, the first 10 PCs are used to generate the t-SNE plot and perform the graph-based clustering.

### GSM models Generation

The scRNA sequences of all the conditions (pre- and post-pneumonia infection in both Wildtype and ZIP8KO conditions) were used to generate the GSM models for macrophage-specific clusters using the ftINIT^34^ method. The gene expression of each macrophage cluster (clusters 1,3,4, 5, 6, and 10) was used to constrain the upper and lower bounds of the reactions based on the gene, protein, and reaction (GPR) associations. By doing so, we obtained GSM models that capture the metabolic capabilities of each of the macrophage clusters distinctly. The same steps were repeated for all the clusters resulting in 24 macrophage models (one model for each cluster in a specific condition). The details of each of the GSM models are available in Supplementary Information.

### Flux Analysis

Flux analysis was conducted of all the macrophage GSM models by using techniques such as Flux Balance Analysis (FBA)^35^, Flux Variability Analysis (FVA)^23^, parsimonious Flux Balance Analysis (pFBA)^36^, and Flux Sum Analysis (FSA)^37^. Each of these methods provides unique insights into different aspects of metabolism, and together, they form a holistic approach to exhaustively characterize the metabolic behavior of each macrophage cluster under all conditions.

### Flux Balance Analysis (FBA), and parsimonious Flux Balance Analysis (pFBA)

FBA provides a single optimal flux distribution under a given condition, usually by maximizing an objective function such as biomass production or ATP generation^35^. pFBA builds on FBA but generates the flux distribution that minimizes total flux while still achieving the optimal objective function value^36^. By integrating both of these approaches, we identified essential pathways for different scenarios such as optimization of energy, or a certain metabolite. In doing so, we identified the pathways, reactions, and metabolites acting as bottlenecks in the ZIP8KO models.

### Flux Variability Analysis (FVA)

FVA is also an extension of FBA that explores all feasible flux distributions by calculating the minimum and maximum for each reaction while maintaining the objective optimally^23^. This ultimately leads to the identification of alternative metabolic pathways that could potentially achieve the same outcome. We used FVA to obtain the flux ranges of each reaction for all the models. This helped us understand the similarities and differences in the metabolic network of each of these clusters for both wildtype (pre- and post-infection) and knockout models (pre-and post-infection)

### Flux Sum Analysis (FSA)

The three above-mentioned methods focus on identifying reaction fluxes that are feasible with our optimal value for the specified objective function. FSA extends the understanding of the metabolic networks at each condition by providing metabolite-level insights. FSA highlights metabolite turnover, bottlenecks, and regulatory points and how they shift from one condition to another^37^. In our case, it helped us understand the metabolic shift and the key metabolites and pathways crucial in eliciting immune response from WT pre-infection to WT post-infection and the bottlenecks due to which the ZIP8KO models were lacking.

### Shadow Price and MetaShiftOptimizer

Shadow Price is a key dual variable obtained from FBA that quantifies the marginal impact of a metabolite’s availability on the objective function^29^. Meaning it helps us identify the key reactions, and metabolites who’s increased or decreased fluxes have a direct impact on the objective. A high shadow price indicates a metabolite is limiting the objective, suggesting that increasing its availability could potentially enhance the objective, and the zero-shadow price implies that the metabolite is non-limiting. Additionally, we also applied a bilevel optimization formulation, MetaShifOptimizer, which allows us to identify key targets, which when manipulated by either upregulation, downregulation, or deletion can help shift the metabolic fluxes to a desired state^30^. For example, we used MetaShiftOptimizer with a knockout pre-infection model as a base and Wildtype pre-infection as a target to obtain the list of potential targets for restoring the immune response. The same method was also repeated for ZIP8KO models’ post-infection to identify targets that could potentially elicit immune functions similar to the wildtype post-infection model. The combination of both of these methods allowed us to identify critical metabolites for both pre-infection and post-infection conditions.

### Isolation and Generation of BMDMs

BMDMs were generated as previously described with a few modifications^55^. Briefly, the femurs and tibias from C57BL/6 WT and *Zip8*KO mice (7-10 weeks old) were harvested, and bone marrow cells were plated in DMEM (cat# 10-014-CM, Corning) supplemented with 10% FBS (cat# 35011CV, Corning), 1% Pen/Strep (cat# 16777164, VWR), and 50 ng/mL recombinant mouse granulocyte monocyte-colony stimulating factor (GM-CSF cat# AF-315-03, PeproTech) at a density of 1.4 x 10^6^ cells/mL in 100 mm dishes.

### Streptococcus pneumoniae Culture

*S. pneumoniae* strain JWV500 (D39hlpA-gfp-Cam’), a generous gift from Dr. Jan-Willem Veening (University of Lausanne, Switzerland), was grown to mid-log phase, aliquoted, frozen, and stored at −80°C until further use. Bacteria were swabbed from frozen stock onto Remel blood agar plates (cat#R01202, ThermoFisher) and incubated at 37°C with 5% CO_2_ overnight. The next day bacteria were grown to log phase in Remel Mueller Hinton Broth (cat# R112478, Fisher Scientific, Lenexa, KS) supplemented with 32 mg/ml chloramphenicol to an OD_540_ of 1.0. For quantification of pneumococci, serial dilutions of the bacteria were plated on Remel blood agar plates, incubated at 37°C with 5% CO_2_ overnight to determine colony forming units (CFUs).

### Internalized Bacterial burden

BMDMs were plated at a density of 1×10^6^ cells/mL in 6 well plates and allowed to adhere for 24 hours. Cells were then pre-treated with or without 100nM KYNA (cat#16792, Cayman Chemical), 500µM or 5mM di-ethyl succinate (cat#112402, Sigma-Aldrich), 10µM or 100µM NAC (cat#20261, Cayman Chemical), 10µM or 100µM GABA (cat#A2129, Sigma-Aldrich) for 2 hours in DMEM. Cells were washed with PBS and infected with *S. pneumoniae* (multiplicity of infection of 10) for 60 minutes. Cells were then washed with PBS and treated with 200 µg/mL gentamicin (cat#15750-060, ThermoFisher) and 10 µg/mL penicillin/streptomycin (cat#16777-164, VWR) for 30 minutes to kill the bound and extracellular bacteria. Cells were washed with PBS incubated overnight in fresh BMDM media with or without KYNA, NAC, succinate or GABA at designated concentrations. Cells were washed with ice-cold PBS and lysed with water. Serial dilutions of lysates were plated on blood agar plates and following overnight incubation colony forming units (CFUs) were enumerated.

### ROS Detection

BMDMs were plated at a density of 1×10^5^ cells/well in a black 96-well plated and were allowed to adhere overnight. Cells were treated with 5µM MSR (cat#M36008, ThermoFisher) in HBSS for 30 minutes at 37°C, washed then infected with *S. pneumoniae* with a MOI of 10:1 for 60 minutes. Fluorescence was quantified using a plate reader (SpectraMax i3x) at excitation/emission 540/590 nm per manufacture’s protocol.

### Bioenergetic Assessment

Glycolytic and mitochondrial stress tests were performed using the XFp Extracellular Flux Analyzer (Agilent). BMDMs were plated at a density of 1×10^5^ cells/well in 8-well Seahorse XFp cell culture miniplate. Plates were centrifuged at 300 x g for 5 minutes to ensure cells were seeded at the bottom of the well then incubated at 37°C in 5% CO_2_ overnight. Cells were washed and incubated in fresh DMEM or infected with *S. pneumoniae* at a MOI of 10:1 in DMEM for 1 hour at 37°C at 5% CO_2_. Following infection, cells were washed and equilibrated in Seahorse XF DMEM medium, pH 7.4 (cat# 103575-100, Agilent) supplemented with 10 mM glucose, 1 mM sodium pyruvate, and 2 mM L-glutamine, and incubated at 37°C without CO_2_. The mitochondrial function was assessed according to the manufacturer’s protocol for the Seahorse XFp Mitochondrial Stress Test. Oxygen consumption rate (OCR) and extracellular acidification rate (ECAR) were measured over time following a series of injections with oligomycin (1.5 uM), carbonyl cyanide-4-(trifluoromethoxy) phenylhydrazone (FCCP) (1.0 uM), and rotenone/antimycin A (0.5 uM). Glycolytic function was measured using the protocol for the Seahorse XFp Glycolysis Stress Test. ECAR was measured over time following a series of injections with glucose (10 mM), oligomycin (1.0 uM), and 2-deoxyglucose (50mM). Raw values were determined in Seahorse Analytics 1.0.0 and normalized to total protein concentration in the sample. Values are expressed as means of biological replicates ± SEM.

### Statistical Analyses

Data were analyzed using GraphPad Prism version 10.2.3 (GraphPad Software, La Jolla, CA). Unpaired Student’s t-test was used to determine differences between groups. Values are expressed as means ± SEM. A value of p < 0.05 was considered significant.

Data were analyzed using GraphPad Prism version 10.5.0 (GraphPad Software, La Jolla, CA). Statistical significance was determined using one-way ANOVA followed by Sidak’s corrections for multiple comparisons between groups. Values are expressed as means ± SEM. A value of p < 0.05 was considered significant.

## Data Availability

ScRNA seq data used in the study is available at request from the corresponding authors. All other data are available in the article and its Supplementary files or from the corresponding author upon request. Source data are provided with this paper.

## Code Availability

The mathematical models generated are available at https://github.com/ssbio/ZIP8_Models

## Acknowledgements

The figures are generated by using Biorender. We would also like to thank the Holland Computing Center (HCC) of the University of Nebraska, which receives support from the Nebraska Research Initiative (United States of America). This research was funded by the National Institute of Health (NIH) R35 MIRA grant (5R35GM143009), awarded to RS, the National Institutes of Health; the National Heart, Lung and Blood Institute Grants # R01-HL156952 awarded to DLK.

## Author Contributions

Conceptualization: SM (Sunayana Malla), RS (Rajib Saha), DLK (Daren L. Knoell), and DRS (Derrick R. Sameulson); methodology: SM, DRS (Daeandra R. Smith), AMP (**Ashley M. Peer**), DLK, DRS, RS; data analysis: SM, RS, DRS; experimental validation: DRS, AP, DLK, DRS; writing – original draft preparation: SM and RS; writing – review and editing: DRS, AP, DLK, DRS, SM, RS; funding acquisition, RS, DRS, DLK. All authors have read and agreed to the published version of the manuscript.

